# Dissecting the control mechanisms for DNA replication and cell division in *E. coli*

**DOI:** 10.1101/308155

**Authors:** Gabriele Micali, Jacopo Grilli, Jacopo Marchi, Matteo Osella, Marco Cosentino Lagomarsino

## Abstract

Understanding how single *E. coli* cells coordinate the timing of cell division with genome replication would unlock a classic problem of biology, and open the way to address cell-cycle progression at the single-cell level. Several recent studies produced new data and proposed different models, based on the hypothesis that replication-segregation is the bottleneck process for cell division. However, due to the apparent contrast in both experimental results and proposed mechanisms, the emerging picture is fragmented and unclear. In this work, we re-evaluate jointly available data and models, and we show that, while each model contains useful insights, none of the proposed models, as well as generalizations based on the same assumptions, correctly describes all the correlation patterns observed in data. This analysis leads us to conclude that the assumption that replication is the bottleneck process for cell division is too restrictive. Instead, we propose that two concurrent cycles responsible for division and initiation of DNA replication together set the time of cell division. This framework correctly captures available data and allows us to select a nearly constant added size per origin between subsequent initiations as the most likely mechanism setting initiation of replication.

## INTRODUCTION

Each cell needs two copies of the genome to divide. The notion that this simple principle must be central for the cell cycle was already clear in early studies [1, 2]. For the model organism *E. coli*, a wealth of information was gathered starting from the late 1950s [3–6], leading to important insights on cell-cycle progression. Today, a relevant set of the key molecules playing a role in the cell cycle of these bacteria is known [7–13]. However, determining exactly how cell division is coordinated with genome replication in *E. coli* is still an open problem. The reason is that our knowledge is still based mostly on population averages, which mask the behavior of single cells [14]. Instead, understanding homeostatic processes in cell-cycle progression needs the knowledge of correlations between subsequent cell-cycle events at the single-cell level. For example, we do not know for sure whether in single cells replication initiation is triggered at a critical size [14–16], whether there are licensing constraints inhibiting initiations [14, 17], and whether the rate-limiting checkpoint for the decision to divide is typically independent from replication initiation or not [18].

A recent wave of experimental and theoretical studies promises to untie this knot, as high throughput single-cell data are becoming routinely available [19–24]. These measurements can in principle access the full correlation pattern between several cell-cycle events, from which we can extract mechanistic interpretations [14]. For instance, there is agreement on the fact that (in most cases) the added volume between consecutive cell divisions is nearly uncorrelated to cell size at birth, a principle sometimes called “adder” [14, 15, 18, 19, 25–28]. However, fundamentally different models that account for this “adder” behavior have been proposed [14–16, 18, 28], leaving us with a complex landscape of models that appear incompatible and in contrast with each other. Additionally, we lack a general theoretical framework to interpret the correlation patterns in the data and to compare and falsify different models.

Here, we aim to provide a solid framework for solving the apparent contradictions between recent claims, by a joint modeling and data analysis approach (considering the available high-quality single-cell data sets). Focusing on the coordination of genome replication with cell division, we first re-analyze currently available data and models. In a parallel study we have introduced the “concurrent cycles” idea: a division-related process (e.g. completion of the septum) and a replication-segregation process (e.g. release of occlusion from the nucleoid) compete for setting cell division [29]. In this study, we develop this idea in two directions. First, we study systematically general models based on the two alternative hypothesis that replication is always or never the limiting process for division. Despite the flexibility and the additional free parameters of these general models compared to the ones proposed in the literature, we show that they irremediably lead to predictions that are inconsistent with available data if the whole pattern of correlations is considered. Second, assuming the concurrent-cycles framework, we ask whether it is possible to capture all the measured correlation patterns and to use them to isolate the specific mechanisms setting initiation and division.

## STAR METHODS

### Method details

#### Data sets

We used published data sets from refs. [15, 21, 25, 28]. The data from refs.[15, 28] contain information on replication initiation and cell division on tracked single cells, and constitute the core of our analysis. Full details on our analysis procedures are provided in SI (section S1). The other data sets only contain growth-division data of tracked single cells and were used for further comparisons of models’ predictions with data.

#### Models

We considered and analyzed different stochastic models for the cell cycle at the single-cell level. These analyses can be divided into three stages: (i) a re-analysis of the published models in refs. [15, 16] (ii) the generalizations and formulations of the models of refs. [15, 16], called “ICD” and “BCD” frameworks, and (iii) the concurrent-cycle model proposed in [29], which has as a limit case the pure adder model advocated by ref. [18]. The full definitions of all the models used here, and our calculations, are presented in SI (sections S4 and S5).

### Quantification and statistical analysis

Coupling parameters between growth in a cell-cycle interval and size (the parameters *λ*_*X*_ in our models) were evaluated from scatter plots as linear fits of binned averages (‘lm’ function in R based on least squares), or alternatively (the parameters 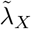) from the equivalent method based on the covariance of two variables (see Mathbox and SI section S4). The two methods are equivalent, but binned averages have the advantage of estimating efficiently the conditional averages defining the control parameters in presence of high noise [30].

The analysis reconstructing the size correlation secondary initiations in the DnaQ data sets from Wallden *et al.* [15] is described in SI, sec. S1 and SI figure S1.

In presence of constraints, i.e., for initiations that were scored in the data only after division, we performed a Bayesian fit of a bivariate Gaussian keeping the constraint into account, and extracted the control strength from the covariance. The algorithm was tested on computational data (full details provided in SI, section S2). Models were analyzed by both direct simulations and analytical calculations.

### Data and software availability

The custom-written code (made of several programs and scripts in C,C++, python and R) generated for statistical analysis and model simulation is available with the authors.

*Further information and requests for resources should be directed to*

## BACKGROUND

### Review of current models of the *E. coli* cell cycle

We start by reviewing currently available models and the key differences in their predictions (Fig. 1) that need to be reconciled. As a premise, all models must reproduce the ubiquitous “near-adder” correlation pattern, robustly found for cell division in several data sets [20], i.e. the fact that added size between consecutive initiations is uncorrelated with initial size (Fig. 1A).

**FIG. 1.**
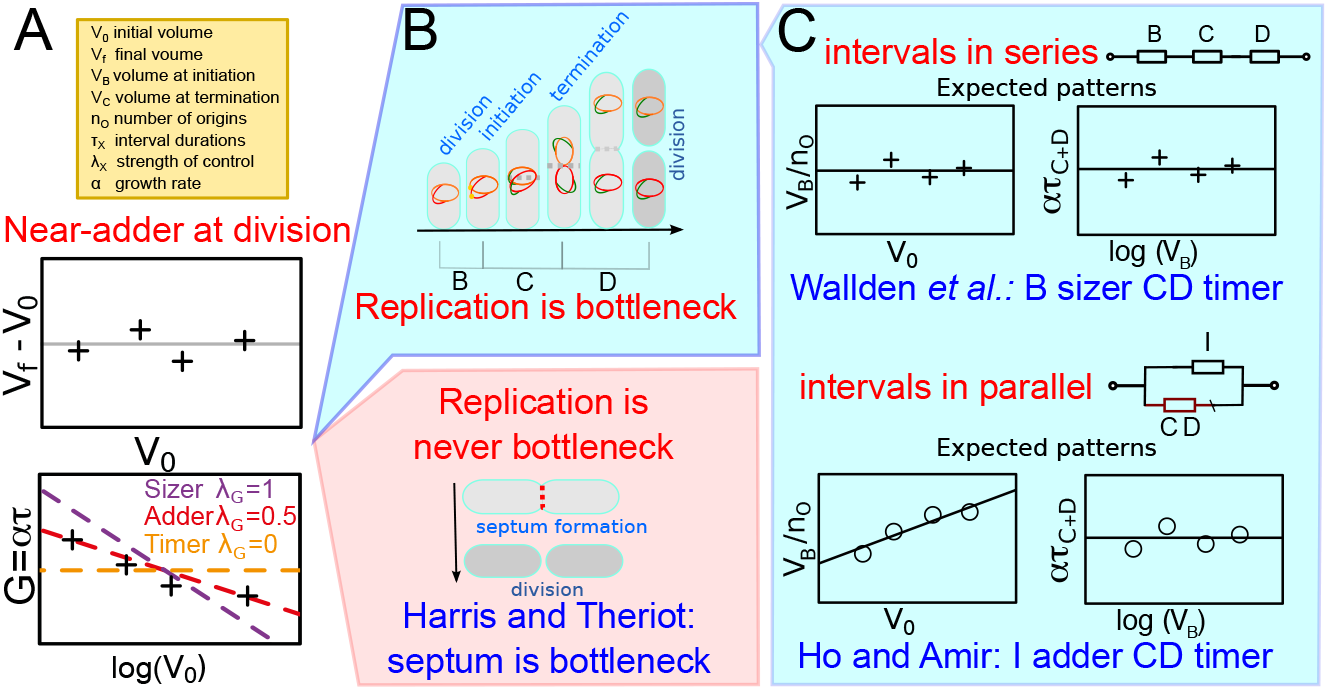
Comparison of the existing models linking cell-cycle progression to cell division in single *E. coli* cells. (A) Top: All models need to comply to the robust “near-adder” pattern found in data (added size over the cell cycle uncorrelated with initial size). Bottom: The near-adder pattern in the “size-growth” plot of net growth *G* = log(*V*_*f*_ */V*_0_) = *ατ* vs logarithmic initial size log(*V*_0_) has a negative slope 1/2. The slope parameter *λ*_*G*_ between 0 (no control) and 1 (absolute threshold) quantifies control strength. (B) Opposite hypotheses for cell division: (top) completion of replication and segregation is always rate-limiting [15, 16], defining the “*B, C, D*” periods, or (bottom) it is never limiting, and division may be triggered by, e.g. a threshold amount of surface material necessary to form the septum [18]. (C) Models in which replication/segregation is bottleneck. The cartoon plots summarize the expectation, in each model, for the expected correlation patterns between the volume at birth, *V*_0_ and the size at initiation per origin *V*_*B*_*/n*_*O*_ (left plot) and between *ατ*_*C*+*D*_ and the logarithmic initiation size log *V*_*B*_, where *τ*_*C*+*D*_ is the period between replication initiation and division (right plot). Wallden and coworkers [15] (top) assume that the periods associated to replication and segregation are consecutive and juxtaposed “in series”. This model postulates a critical size per origin at initiation, and a duration of *τ*_*C*+*D*_ that is coupled to single-cell growth rate but not to cell size. In the Ho-Amir model [16] (bottom), the timing between initiation and division *τ*_*C*+*D*_ and between subsequent initiations, *τ*_*I*_ run “in parallel” from a single initiation event. Only *τ*_*I*_ is coupled to size in a way that a constant size per origin is added between successive initiations.

Harris and Theriot [18] assume (Fig. 1B) that the process of replication and segregation is never the bottleneck process for cell division, since it is typically completed well in advance, before other rate-limiting processes trigger division. They also propose that, of the multiple checkpoints needed for cells to complete division, the one that is rate limiting could be the accumulation of a target surface material (enough to build the septum). Under this assumption, and further assuming that the surface synthesis rate is proportional to cell volume, one finds near-adder correlations (SI text, section S3).

Clearly, a main assumption of this model is that genome replication/segregation is typically faster than the critical accumulation of the factor triggering division, and thus that cell division can be unaware of the chromosomes. The studies measuring how different perturbations affect mean cell size conclude that this is not generally the case [31, 32], but the work by Harris and Theriot does not address the evidence pointing to a link between replication and cell division [6, 13]. Thus, in the best-case scenario, it needs to be complemented with a description of DNA replication.

The prevalent view, assumed by all other available models, is that instead the bottleneck process for division is the completion of replication and segregation. We focus specifically on the link between replication and cell division (Fig. 1BC). In this case the cell cycle is naturally divided into the *B, C, D* periods defined by replication initiation, the duration of replication, and cell division, (Fig. 1B), and analogous to the G1, S, G2/M periods of eukaryotic cell cycles.

In particular, there are two main models that are used to explain cell division based on the idea that the replication/segregation process is rate-limiting. Both models are based on the assumption that cell division takes place at a size-independent time after initiation of replication, but they differ in how replication initiation is controlled. A model by Ho and Amir [16] assumes that cells add a constant (i.e., independent from the cell size at initiation) size per origin between initiations (adder between initiations), while division is set by a constant time *C* + *D* (timer) after initiation. A competing model by Wallden and coworkers [15] proposes that initiation is triggered by a constant cell size per origin. This “sizer per origin” extends to single cells the classic picture [6, 12] that assumes a sizer at initiation, motivated by population-averaged data [14]. Both models are compatible with this constant *average* size per origin at initiation, thus with empirical observations at the population level. However, from a single-cell perspective the assumption of a critical size per origin is radically different from the assumption of a constant added mass since the last initiation event. It is an open question whether and how the available data allow to distinguish between these two alternative scenarios.

### Invalidation of current models based on the C+D correlation patterns

We have shown in a parallel study [29] that the existing models cannot capture correlation patterns in the *C* + *D* period. This section recapitulates these inconsistencies between data and model predictions in more detail. For each model describing the replication-division cycle, Fig. 1C shows two key predicted correlation patterns for the *B* and *C* + *D* period. First, the volume per origin at initiation *V*_*B*_ versus initial volume *V*_0_, testing the existence of a size threshold for initiation in single cells. This plot has slope zero if a size-threshold exists (a “sizer”). Second, the correlation pattern between the growth in the *C* + *D* period, *ατ*_*C*+*D*_ and the initial size, testing a possible coupling between the replication-segregation cell-cycle interval and cell size. The slope of this plot (*λ*_*C*+*D*_) is zero if the *C* + *D* period is uncoupled to size (a “timer”).

The two contrasting models by Ho-Amir and by Wallden and coworkers agree in the claim that there is no control between initiation and division, *λ*_*C*+*D*_ = 0 (Fig. 1C), but the duration of this period fluctuates around a cell-size independent value (a timer). A third study by Adiciptaningrum and coworkers [28], found experimentally that the duration of the “D period”, the time between termination of replication and cell division was anti-correlated with size at termination of replication. This coupling between *D* period duration and cell size is at odds with the assumptions of both models. This pattern is confirmed by data from Wallden and coworkers, as shown in Fig. 2, and the existing models based on replication/segregation as the rate-limiting process do not reproduce it. Additionally the assumption that DNA replication is never a bottleneck as in Harris-Theriot model [18] leads to quantitatively wrong predictions in the single-cell correlations patterns of the replication-related cell-cycle intervals. We will discuss in more detail this question in the following, as this model represents a specific limiting case of the concurrent-cycles framework.

**FIG. 2.**
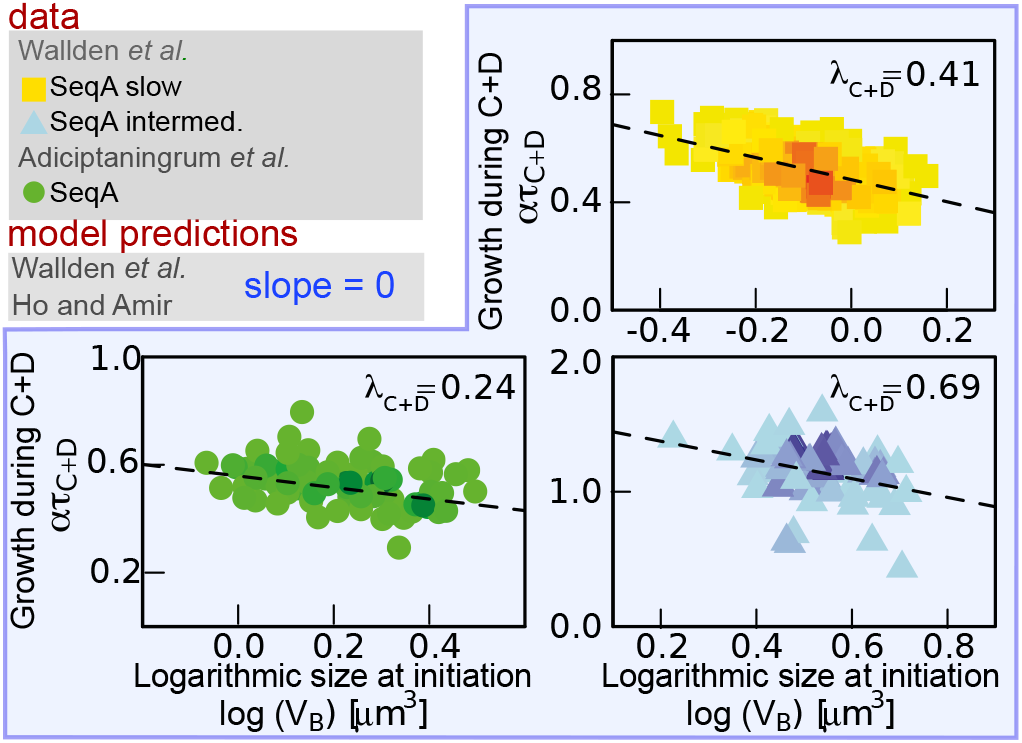
Current models fail to capture the experimental correlation patterns of the cell-cycle intervals related to replication and segregation (*C* + *D* period). Scatter plot and slope of the size-growth plot for the *C* + *D* period, i.e. *ατ*_*C*+*D*_ as a function of initiation size log(*V*_*B*_). Data from ref. [15], slow (yellow squares) and intermediate (light blue triangles) growth conditions, and from ref. [28] (green circles). All data sets presented were obtained by labelling SeqA molecules. The negative values of the slopes (dashed lines from linear fits of binned data), quantifying the parameters *λ*_*C*+*D*_, robustly show size-coupled growth during the *C* + *D* period.

To further explore the limitations of the classic hypothesis that division is limited by replication and segregation, we introduce two classes of models that generalize the two descriptions by Ho-Amir and by Wallden and coworkers. These generalizations show that the assumption itself of replication/segregation as the single rate-limiting-process for division leads to predictions that cannot be reconciled with empirical data, even when the correlation pattern in Fig. 2 is captured with *ad hoc* ingredients.

## RESULTS

### Definition of generalized models based on replication and segregation as the rate-limiting process for division

We now define a general modeling framework that assumes that replication-segregation bottlenecks cell division, with the scope of highlighting the limitations of this assumption. The two models by Ho-Amir and by Wallden and coworkers fundamentally differ in the assumption of how the cell-cycle intervals corresponding to the *B, C, D* periods are temporally juxtaposed (see the sketches in Fig. 1C). In the model by Wallden and coworkers, the cell-cycle intervals are placed “in series”, and no interval can begin if the previous one is not completed. Conversely, in the Ho-Amir model, there is an overarching interval connecting subsequent initiations, and the interval corresponding to the *C* + *D* period runs “in parallel”. Thus, the *B* period in this model is a result of the two parallel cell-cycle intervals (their difference, in absence of overlapping replication rounds).

Therefore, we consider two general “wiring diagrams” of couplings between size and growth (Fig. 3) which we call “ICD” (in parallel) and “BCD” (in series). In these general schemes, each cell-cycle interval is drawn as an arrow, and size is coupled to duration of the interval by generic parameters. These parameters can represent controls acting in case of a size fluctuation, e.g. by reducing the duration of the period in case the cell is larger than average at entry. Each cell-cycle interval is characterized by the coupling parameter *λ*_*X*_ between its duration and cell size (e.g., volume). Such parameters are evaluated directly from the “size-growth” plots of relative growth during each interval vs initial size (illustrated in Fig. 1A) and may range from 0 (timer, no control) to 1 (sizer, absolute size threshold).

**FIG. 3.**
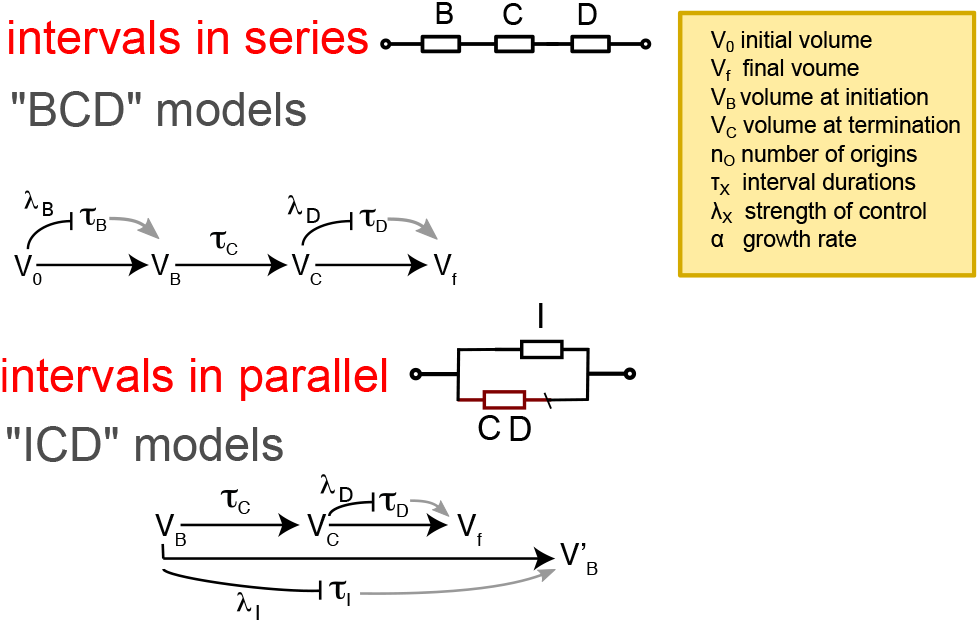
Scheme of the generalized models BCD and ICD. General schemes of models in series (“BCD”) [28] and in parallel (“ICD”). In the sketches, each model is characterized by the control parameters *λ*_*X*_ coupling the growth during a cell-cycle interval and cell size. Noise parameters in each model describe the variability of each cell-cycle interval at fixed initial size.

Specifically, we aim to show that in models assuming that replication is always bottleneck, even parametric generalizations, which are in principle more flexible [28], still lead to predictions that deviate from data.

### Indecisive evidence for a critical size threshold at replication initiation from direct measurements

Before addressing the models, we need to review the experimental support in single cells for the assumption of a sizer at initiation. Clear evidence for a size threshold at initiation would limit the parameter space of our generalized models. Therefore, we re-analyzed the available data in order to test the correlation patterns in the *B* period and to compare them to model predictions or assumptions. Fig. 4 summarizes the results of this analysis.

**FIG. 4.**
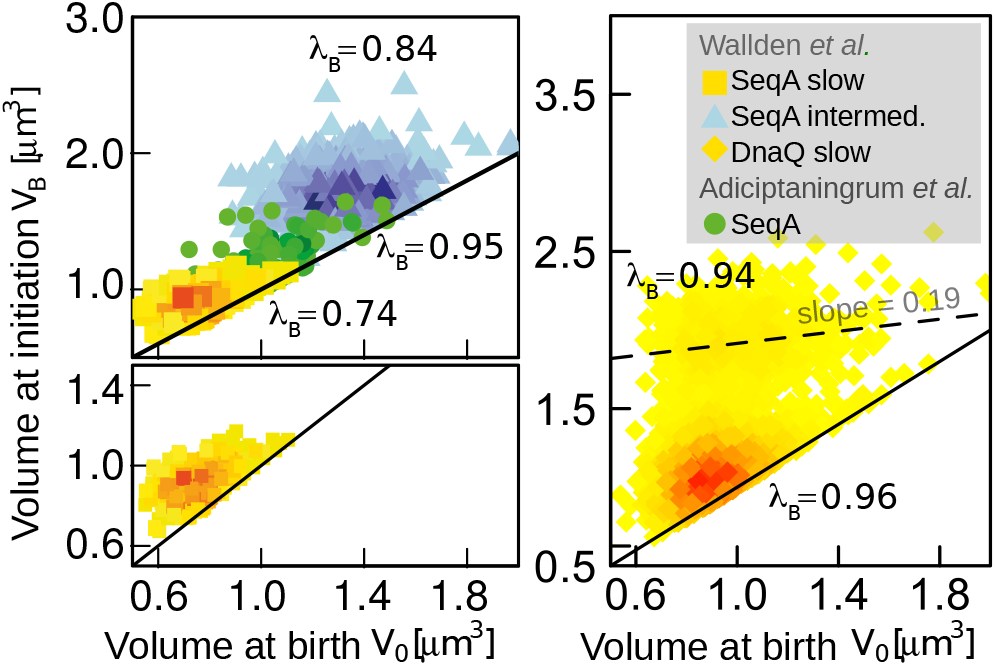
Inconclusive empirical evidence for a sizer at initiation. Volume at initiation vs birth volume (top-left panel). SeqA data from ref. [15], slow (yellow squares) and intermediate (light blue triangles) growth conditions, SeqA data from ref. [28] (green circles). Color scales correspond to the probability density (see SI text). The solid line have slope one (*V*_*B*_ = *V*_0_). Comparison of the same plot between SeqA (left) and DnaQ (right) data from ref. [15], slow growth conditions. Data from DnaQ show the possibility of a second initiation per cell cycle (see SI section S1). Conversely, SeqA foci are detected only after division [14]. The weak dependency of the size of the second initiation with initial size (dashed line is slope from binned average) is only loosely consistent with a sizer (secondary initiations are extracted from foci subcellular localization, tracking, and cell size, Fig. S1). The annotated values of *λ*_*B*_ are extracted from the equivalent size-growth plots (Fig. S2) by Bayesian fits keeping into accounts the constraints.

As noted previously [14, 15], data on replication initiation obtained by labelling SeqA molecules show a constraint whereby foci appearance is recorded only when *V*_*B*_ > *V*_0_. This constraint is visible in Fig. 4 as a “cut” in the correlation clouds between initial volume *V*_0_ and volume at initiation *V*_*B*_. The cut cloud is due to unrecorded initiation events in the previous cell cycle (which is typically not available or tracked in the data set). In order to estimate the slopes *in presence* of the constraint, we performed a Bayesian fit of a bivariate Gaussian using the data in Fig. 4 under the assumption that the data below the constraint were censored (see SI section S2 for the description and testing of this algorithm). The results are not far from a sizer, but also deviate sensibly from this pattern.

DnaQ foci, instead, are more visible before division, although these data are affected by false-positive detection due to blinking and by poor segmentation-tracking of cells [15]. Despite of these problems, the DnaQ data show evidence of double initiations in the same cell cycle and allow a different analysis, since the data of secondary initiations are free from the cut in the correlation cloud. Secondary initiations can occur in cells with delayed divisions (which might meet the size criteria for initiation twice in the same cell cycle). We have performed a refined analysis of the *B* period in the DnaQ data, defining secondary initiations from information on foci subcellular localization and time tracks of both foci and cell size (see SI sec S1 and SI Fig. S1).

Fig. 4 shows the cell volume at initiation (*V*_*B*_) against the cell volume at birth *V*_0_ (not shown in the Wallden *et al.* study). We conclude that (i) at the same range of volumes at birth, a secondary cloud of initiations at large volumes appears for DnaQ. [33] (ii) The correlation in the secondary cloud of Fig. 4 confirms the idea that initiation size is weakly correlated with birth size, generally close but slightly divergent from the pattern expected from a sizer. Note however that, because of blinking, we consider DnaQ data less reliable than SeqA for the primary cloud of initiations. This is because mother-daughter progression is not tracked, and as a consequence it is not possible to reliably assign initiation to the first cloud, since the appearance of foci early in the cell cycle could be due to a a real initiation or to a blinking event in the mother cell.

In conclusion, the available direct measurements of initiation size do not conclusively point to the presence of a size threshold at initiation, and generally show a weak but noticeable positive correlation of initiation size with birth size.

### Inconsistencies between the measured correlation patterns for the B and C+D period and generalized models

Having reviewed the main experimental correlations, we go back to the generalized models of Fig 3 to analyze if they may reproduce them. Specifically, we asked whether the observed correlation patterns in C+D (Fig. 2) and in the B period (Fig. 4) could be reproduced jointly and consistently by these models (SI section S4 and S5 and Fig. S4).

A non-zero control variable *λ*_*C*+*D*_ coupling initiation size to division (as the one that is built in the Adiciptaningrum *et al.* model) can be added in a straightforward way to any model. This would trivially make the models able to reproduce the correlation trends in C+D (Fig. 2), although this extra parameter does not have a natural interpretation.

We then considered the *B* period. An issue raised by Fig. 4 is the question of how adder correlations between divisions can be compatible with a scenario where initial volume and initiation volume are at most weakly correlated. In particular, if we assume an adder between initiations, we need to explain the weak correlation between initiation size and birth size. However, we found that in presence of noise the ICD model (and thus the Ho and Amir model as a particular case) can predict low correlations between initial size and initiation size (hence high *λ*_*B*_, see SI section S5 and SI Fig. S4). Therefore, the correlation pattern in the *B* period (Fig. 4) is not sufficient to distinguish between the models and one has to consider other observables (see SI Section S6 and Fig. S5).

The failure of the entire framework emerges when the patterns for the *B* and *C* + *D* periods are considered jointly. To further test whether the ICD and BCD models could be consistent with data, we solved them analytically, obtaining consistency relationships between the different control parameters. The main relationships are shown in Mathbox. These mathematical expressions are valid in the approximation where the number of overlapping rounds does not fluctuate, but they agree well with simulations (SI Figure S6). While in both ICD and BCD models one is allowed to tune the control parameter *λ*_*C*+*D*_ between replication initiation and the corresponding division event, both models have to follow a general relationship between the inter-division control strength *λ*_*G*_ (see Fig. 1A) and the product of the *B*-period and *C* + *D*-period coupling parameters, (1−*λ*_*G*_)^*n*^ = (1−*λ*_*B*_)·(1−*λ*_*C*+*D*_), where *n* is the number of overlapping replication rounds (see Mathbox, SI section S6, and Fig. S6). Additionally, this relationship does not depend on the noise levels, differently from the one relating the inter-division and the inter-initiation patterns (Mathbox). Thus, it is expected to be robust in the data. This relationship is verified in simulations (SI Fig. S6), but Fig. 5 indicates that the data are in disagreement. This analysis leads us to suggest that the strict assumption that replication is the sole bottleneck for cell division leads to inconsistencies with data.

**Figure 5.**
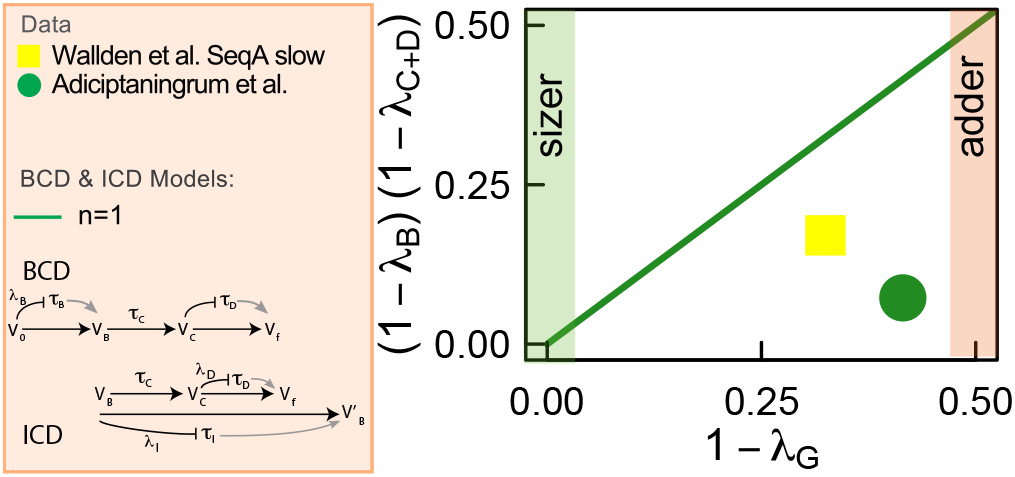
Discrepancy of data with general models assuming that replication-segregation is the bottleneck process for cell division. The plot tests the relationship between predicted size-control between consecutive divisions and the observed correlation patterns for the *B* and *C* + *D* periods. For both the BCD and ICD models (see Mathbox), this is predicted to be an identity in absence of overlapping rounds. This prediction is matched well by simulations (SI Fig. S6), but is violated in data, for the two available conditions in absence of overlapping rounds, casting doubts on both models, even in their parameter-flexible formulations.

### “Sizer theorem”: If any subperiod is in series with a sizer, interdivision time is a sizer

To fully show that existing models based on the hypothesis that replication-segregation is the limiting step for cell division fail, we need to deal separately with the model proposed by Wallden and coworkers [15], which is based on some extra specific assumptions that transcend the ICD/BCD framework. One of the main reasons for the failure of the ICD and BCD frameworks shown in Fig. 5, is that the strength of the control of the *B* period affects the control between divisions. This could be termed the “sizer theorem”: a near-sizer at initiation (witnessed by weak correlation between initial size and initiation size), as well as in any cell-cycle interval in the chain of events leading to cell division, leads to near-sizer correlations between subsequent divisions. This fact, shown by simulations in Fig. 6A, is simple to derive theoretically, and is a consequence of the fact that once cell-cycle progression hits a sizer (hence a trigger that is uncorrelated to the initial size of the previous subperiod), memory of all previous sizes is lost, hence the division size will be uncorrelated to the initial size [27].

**Figure 6.**
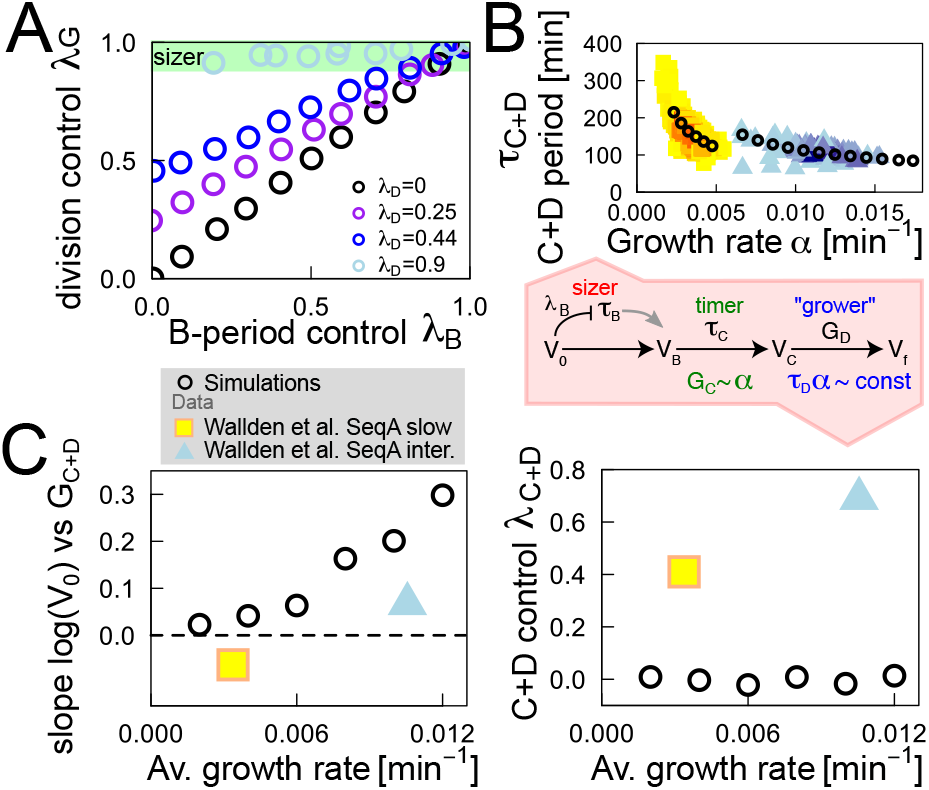
Mother-daughter correlations bypass the sizer theorem, but the resulting model is still in contrast with available data. (A) “Sizer theorem”: if any cell-cycle interval is a sizer, and intervals are placed in series, the interdivision correlation pattern is a sizer. The division cycle control *λ*_*G*_ is plotted as a function of the growth control parameter *λ*_*B*_ for the *B* period, in the BCD model. The division control increases with increasing control over the *B* period and *D* period (*λ*_*D*_ = 0 black, *λ*_*D*_ = 0.25 purple, *λ*_*D*_ = 0.44 blue, *λ*_*D*_ = 0.9 light blue circles). In particular, a sizer at replication initiation implies a sizer on the division cycle, regardless the level of control on the *D* period. (B) Model in which the *B* period is a sizer, *C* is a timer and *D* is a “grower” (see main text), similar to the model in ref. [15]. This model incorporates the observed trend between individual-cell growth rate and C+D period duration as a consequence of the timer+grower pattern. (C) The resulting model is at odds with the Wallden *et al.* data. The correlation between the logarithmic size at birth and the growth during the *C* + *D* period (left panel) is predicted to be positive (circles correspond to simulations), but is close to zero in the data (filled symbols) from ref. [15]. Conversely (right panel), The size control parameter during the *C* + *D* periods is predicted to be zero (circles) while experimental data clearly deviate from this value (filled symbols). Parameters: In panel A, the simulations are performed varying *λ*_*B*_ while keeping fixed the noise ratio *σ*_*C*+*D*_*/σ*_*B*+*C*+*D*_ = 0.7. The other parameters are inferred from data of Adiciptaningrum et al [28]: *σ* = 0.0053 min^−1^, 〈*t*_*B*_〉 = 30 min, 〈*t*_*C*_〉 = 78 min, 〈*q*_0_〉 = 0.03, 〈*n*〉 = 1. In panels B, C and D, the average growth rates range from 0.002 to 0.012 min^−1^ with constant CV equal to 0.2. Mother-daughter correlation of growth rate is set at *ρ* = 0.5. In this modified Wallden model, the average C period is set to 42 minutes with CV 0.1, and for the grower period *ατ*_*D*_ is set to be 0.6 on average, with CV 0.1. The experimental values for the Wallden *et al.* slow growth condition (yellow squares) are ≈ −0.06 and 0.41 for the C and D panels, respectively. For the Wallden *et al.* intermediate growth condition (light blue triangles) are ≈ 0.06 and 0.69 for the C and D panels, respectively. The average size per origin at initiation is set to *ν* = 0.9 [*μ*m^3^] with constant CV of 0.1 for all panels.

Therefore, any model assuming a sizer at initiation has to bypass the sizer theorem in order to be compatible with near-adder correlations between subsequent divisions.

We now proceed to show how the model proposed by Wallden and coworkers solves this problem, but we will also show that this solution leads to predictions that are falsified by the available data.

### The Wallden *et al.* model escapes the sizer theorem thanks to correlations across generations and stochasticity of single-cell growth rates

Wallden and coworkers [15] simulate a model where initiation is triggered by a sizer, compatible with our re-analysis of their experimental data and Fig. 4. However, the interdivision correlations in this model are not compatible with a sizer, violating the sizer theorem (Fig. 6A). To explain how this is possible, the authors argue with simulations that the result is due to a direct growth-rate dependency of *C* + *D* period duration to growth rate (Fig. 6B), which they fit empirically from data with a power law. The direct link between this feature and the model behavior is not clarified in their work, nor is the reason of the near-inverse relationship between the single-cell growth rate and the duration of the *C* + *D* period, which is taken as an empirical fact.

In order to shed light into this result, we introduce an equivalent parameter-poor model, where a simpler ingredient leads to the same behavior without the need for arbitrary phenomenological fitting procedures. Specifically, we assume (Fig. 6B) that (i) initiation is driven by a sizer as in the standard version of the model, (ii) the *C* period is a timer and (iii) the *D* period is a “grower”, i.e. the net growth quantified by *αD* = log(*V*_*f*_ */V*_*D*_) is uncor-related to size at replication termination log(*V*_*D*_). This last ingredient is different from a simple timing mechanism. It implies that, e.g., in cells where is larger, the *D* period duration will tend to be shorter, hence the two variables in the product *G*_*D*_ = *ατ*_*D*_ become naturally anticorrelated. Conversely, in the case it were a timer, the net growth would be still uncoupled to initial size, but the fluctuations of the subperiod duration would not be coupled to single-cell growth rate.

The assumed “grower” correlations for the *D* period lead to define a model that behaves equivalently to the model of Wallden and coworkers, with the advantage that the coupling of *C* + *D* period duration with growth rate is not adjusted by hand, but is a natural consequence of the “grower” assumption. We were able to solve this model analytically for the inter-division correlation patterns, revealing the explanation of the effect found by Wallden and coworkers, and we compared it with simulations and empirical data.

Our prediction is that growth-rate correlations (quantified by Pearson correlation *ρ*) across generations give rise to a correlation loss in the inter-division size-growth plot with respect to the sizer. We also predict that an analogous effect is expected in presence of overlapping replication rounds because of the correlations induced by a *C* period lasting over multiple generations (SI Figure S3). The mechanism is the following. A cell that grows at faster rate than average will typically divide at a larger size. The growth rate of the subsequent generation will thus retain a memory of initial size (through its correlation with the growth rate of the previous generation). Since the *C* + *D* period is partly relying on a timer and partly on a grower, its anti-correlation with the growth rate will create an effective correlation of its duration with *initial* (and not initiation) size and hence weaken the sizer correlation between divisions (see SI text for a full explanation and calculation).

This prediction is in line with the arguments provided by Wallden and coworkers. It is also in excellent agreement with simulations, but the agreement is not very satisfactory when compared to empirical data from several published studies of inter-division correlation patterns (SI Figure S3).

Additionally, and crucially, these ingredients do not produce the correct correlation patterns for the *C* + *D* period (Fig. 6C). Indeed, the Wallden *et al.* model, as well as our variant (which is equivalent), predict that there should be (negative) correlation between the *initial* cell size and the growth during the *C* + *D* period, but no correlation between *initiation* size and growth in the *C* + *D* period. The former is a consequence of the memory effect carried by the persistence of the individual cell growth rate and the timer+grower pattern of the *C* + *D* period, while the latter is a consequence of loss of memory of initial size at initiation given by the sizer mechanism (sizer theorem). Fig. 6C shows that empirical data from the Wallden *et al.* study follow exactly the opposite pattern, and show stronger correlations between *C* + *D* period duration and initiation size, and weaker correlations between *C* + *D* period duration and birth size.

Thus, while we have provided a simple rationale for the model proposed by Wallden and coworkers, and we support the observations that lead to its definition, we can also conclude that the basic ingredients of this model cannot be fully correct.

### A concurrent cycles model based on competition between adders explains the correlation patterns from simple ingredients

All the above considerations give a fairly complete account of the problems encountered by trying to explain available data with models that assume that replication/segregation is always bottleneck for cell division. In particular, these models fail to capture crucial features of the *C* + *D* period (Fig 2) or the composite pattern of correlations in *B*, *C*+*D* and the overall cell cycle (Fig. 5).

We now turn to use this knowledge to pinpoint the size-regulatory processes in the concurrent-cycles framework [29]. In this model the competition between two concurrent processes, one setting division and one controlling the initiation of DNA replication, naturally reproduces the pattern in Fig. 2 and in Fig. 6B without ad hoc assumptions or additional parameters [29]. The basic idea is to relax the hypothesis that replication-segregation is always the bottleneck process and assume that concurrent cycles regulate division and the replication-segregation cycle. In other words, the model assumes that neither the limit where the bottleneck process is always chromosome segregation [15, 16, 28], nor the limit where segregation typically finishes before division [18] are realized.

As we have shown, empirical data leave open the question of the control of replication initiation (Fig. 4). Hence, we used the concurrent cycles framework in combination with the available data to pinpoint the most likely mechanism setting replication initiation. Following Harris and Theriot (and as suggested by several empirical observations, see Fig. 1A), we started by assuming that the interdivision process is a near-adder. We then compared two variants where initiation is set by a sizer per origin, or by an adder per origin between subsequent initiation events.

After each initiation, a minimum (size-uncoupled) *C* + *D*′ period is necessary before division. *D*′ may not be the observed duration of the *D* period in the case in which the completion of the replication-segregation period is not the bottleneck event for cell division. The slowest of the two concurrent processes decides division (Fig. 7A).

**Figure 7.**
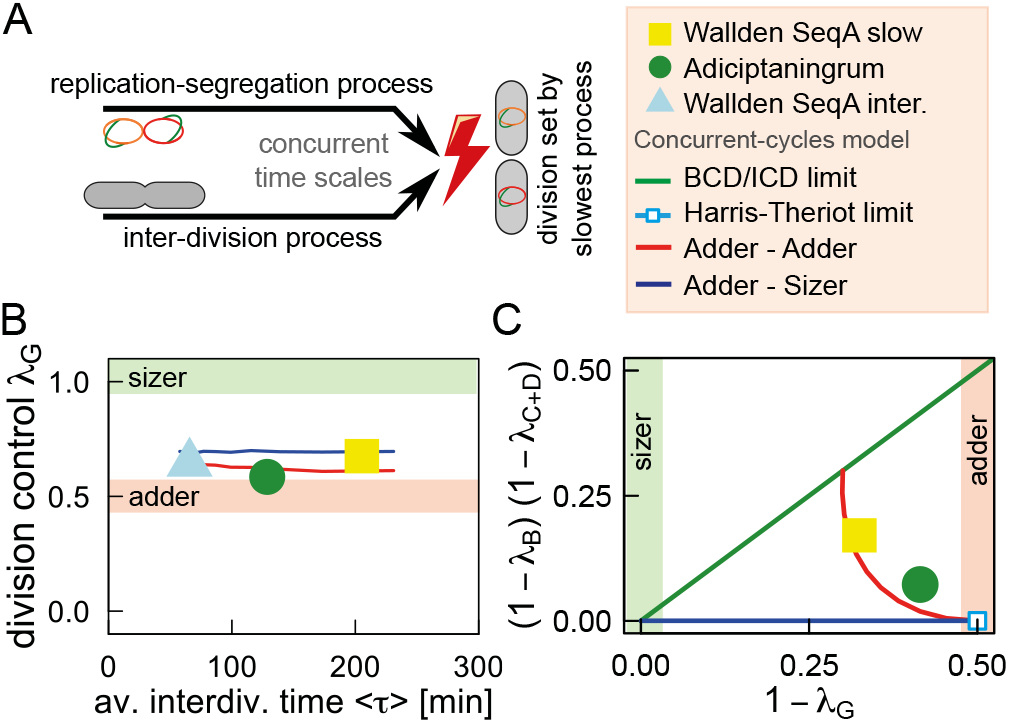
The correlation patterns of the concurrent-cycle model fully agree with available data and support the hypothesis of a near-adder per origin between-initiations. (A) Schematic of the concurrent-cycles model: an inter-division process and a process setting replication initiation compete for the decision of the cell to divide. We considered the cases where an adder between divisions concurs with an adder between initiations (red lines) or a sizer (blue lines) at initiation. (B) The division control parameter *λ*_*G*_ as a function of the average interdivision time gives a near-adder for both concurrent models, in agreement with data. The predictions assume that there is not a clear separation of the size scales of the two processes (*p*_*H*_ ~ 0.5), thus that competition between processes is actually present. (C) Comparison of empirical data (without overlapping replication rounds) with the predictions of the adder-adder vs sizer-adder models allows to select the mechanism setting initiation. The green lines correspond to models based on replication-segregation as the single rate-limiting process for cell division (BCD and ICD models). The limit case (light blue, corresponding to the Harris and Theriot hypothesis or *p*_*H*_ ≃ 1) of a chromosome-agnostic division also fails to reproduce the data. Only the concurrent-cycles model with an adder between consecutive initiations and an inter-division adder (red line corresponding to a noise ratio 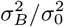 fixed from data) can match the data, by varying the only free parameter *p*_*H*_ (the probability that replication is not the bottleneck in a cell cycle) in a relatively narrow range around 0.5 (*p*_*H*_ = [0.25, 0.75]), compatible with our assumption of competition between two processes. Numerical simulations and data with overlapping replication rounds are shown in SI Fig. S7.

An important quantity in this model is the probability *p*_*H*_ that the inter-division process is the limiting one. This parameter is an output of the model, and depends on both concurrent processes. It is related to the typical added size between divisions and its variability for the inter-division process, and the typical size at initiation and duration of the *C* + *D*′ period in the replication-related cycle.

Importantly, the concurrent cycle framework only relies on parameters that are fixed from data or from natural assumptions, and not forcedly adjusted. The control strengths of the two concurrent processes *λ*_*I*_ and *λ*_*H*_ are fixed *a priori* by the specific assumed mechanisms for the concurrent cycles, as in previous models (we considered the cases of adder-adder and sizer adder, see Mathbox). Importantly, there is an additional time scale (or equivalently a size scale) that needs to be defined in this framework. Specifically, this time scale is captured by the parameter *D*′, and depending on its relation to the natural time scale of the system set by the doubling time, it will define which of the two processes is more likely to be the slowest (i.e., the value of *p*_*H*_). Equivalently, the two size scales associated to the processes correspond to the mean size per origin at initiation, which several studies indicate is remarkably constant in different conditions [31, 32], and to the average added size. However, in order to have the two concurrent processes to be actually in competition, their associated time or size scales have to be comparable, and this condition fixes a narrow range for the parameter *D*′, and consequently for *p*_*H*_.

Under the assumption that the two time (or size) scales are comparable, both model variants have a comparable number of parameters (or less) as the Ho-Amir and Wallden *et al.* models, and robustly give a near-adder correlation pattern across divisions (Fig. 7B), with a slight deviation from the “pure” adder prediction (*λ*_*G*_ = 0.5) which is compatible with the data. Additionally, we have shown [29] that both model variants naturally capture the correlation patterns relative to the *C* + *D* period (Fig. 2 and Fig 6B) without the need of adding these trends as *a priori* ingredients.

More interestingly, the model gives different predictions (see Mathbox), depending on the model assumptions, for the *relationship* between the control parameters *λ*_*B*_, *λ*_*C*+*D*_ and *λ*_*G*_. Importantly, these plots allow to test the process regulating replication initiation by the correlation patterns of the concurrent-cycles model (see Mathbox). We find that the data sets coherently support the assumption of an adder per origin between subsequent initiations [16]. Since the control parameters of the model are all fixed by the hypothesis that the inter-division and inter-initiation processes are both near-adders, and the noise parameters can be fixed directly from data, the only parameter that is allowed to vary in these models is *p*_*H*_. We verified that empirical data can be captured by varying this parameter in a relatively narrow range of values around 0.5. This support the idea that the time scales of the two processes are approximately matched and thus competition is typically in place. Moreover, the only two data points obtained with the same strain (the Wallden data sets) fall very close, hence should have a very close value of *p*_*H*_. Additionally, a comparison of these predictions with the empirical data (Fig. 7C and SI Fig. S7) further confirms the limitations of existing models and their generalizations. In particular, the extreme case of replication/segregation as a bottleneck (BCD/ICD limit, corresponding to *p*_*H*_ = 0) fails regardless of the flexibility on the parameters, and the same is true for the opposite limit (Harris and Theriot limit, *p*_*H*_ = 1).

**Mathbox 1.**
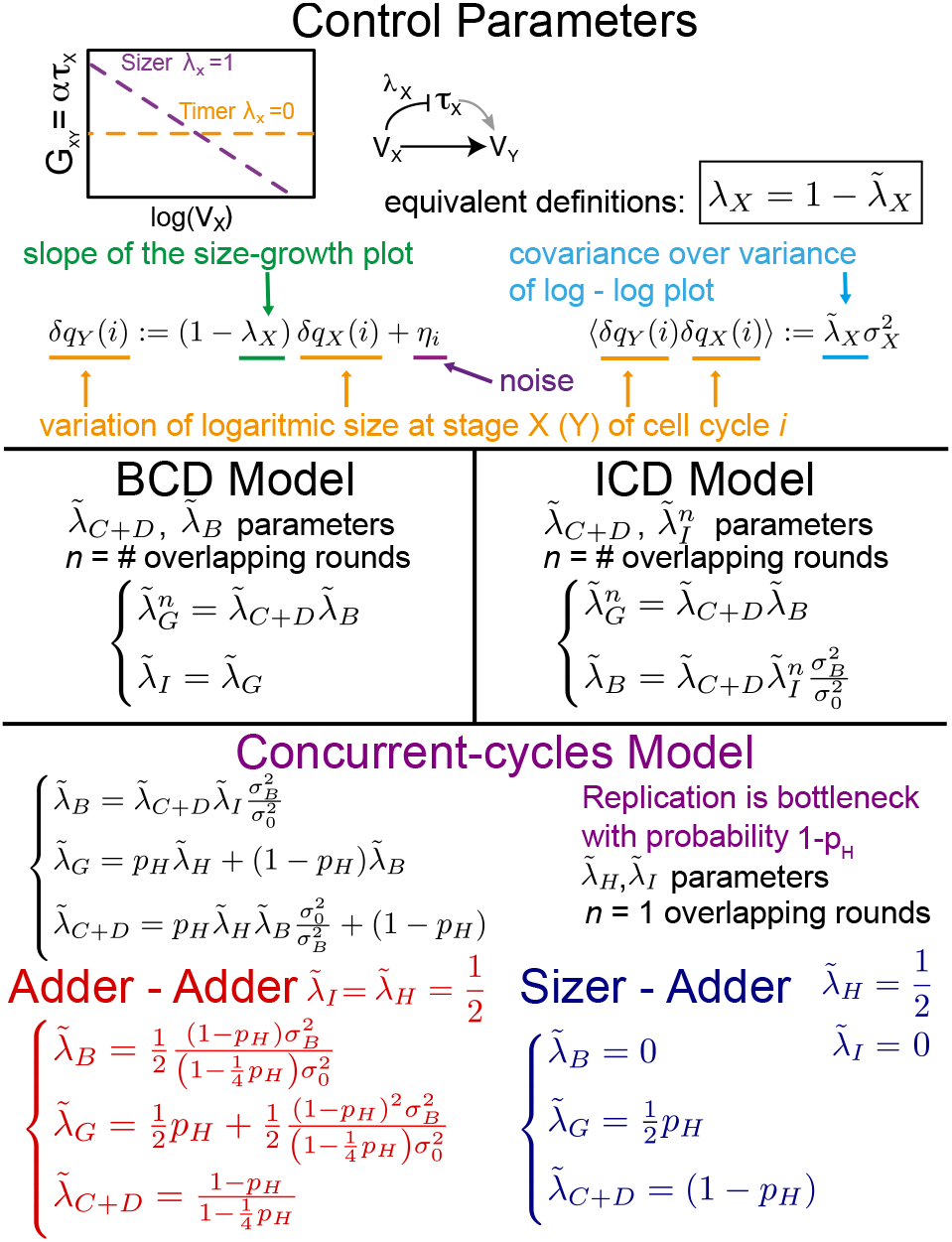
Main relationships between control parameters in the different models. Top panel: definition of control parameters. *δq*_*X*_ = *q*_*X*_ − 〈*q*_*X*_〉 is the deviation from the mean of logarithmic size *q*_*X*_ = log *V*_*X*_ at checkpoint *X* of the cell cycle. The control parameters *λ*_*X*_ (extracted from the slope of size-growth plot as in the sketch) and 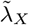 (slope of scatter plot of logarithmic size at the end vs the beginning of the cell-cycle interval) can be used equivalently. Middle panel: main relations between control parameters for the BCD and ICD models. Bottom panel: main relations for the concurrent-cycles model, where *p*_*H*_ is the probability that the inter-division cycle is bottleneck (*p*_*H*_ ≃ 0.6 if the size scales encoded by the two concurrent processes are matched). The red and blue brackets show the particular cases of adder between divisions concurring with adder between initiations and sizer at initiation respectively.

## DISCUSSION

### What is the bottleneck process for cell division?

As clearly explained by previous studies [14, 16, 18] a central point in solving the question of the determinants of cell division in *E. coli* is whether the process of replication followed by segregation is a bottleneck for cell division or not. The more conventional view [15, 16] is that the chromosome cycle is always a bottleneck, and the decision to divide is slaved to the decision to initiate replication through the processes of completing replication itself and segregation. The strongest piece of evidence in this direction are classic and recent observations on mean cell size that we discuss below. The less conventional hypothesis of Harris and Theriot [18], motivated by their measurements of the mean surface/volume dynamics of *E. coli* cells, is that replication is *never* a bottleneck for cell division. Under this assumption, *E. coli* decides to divide independently of the chromosome replication cycle.

Our main result is twofold. First, a wide class of models based on a single rate-limiting process setting cell division is unable to explain at the same time the correlation patterns for the cell-cycle sub-periods. This failure of the replication-based models [15, 16] points to a model where replication is not always bottleneck, but two (or more) processes, of which (at least) one is replication-related, act on similar time scales and compete for setting cell division [29]. Second, assuming this framework of concurrent cycles, our analysis clearly indicates that replication initiation is set by a near-adder per origin between initiations, as suggested by Ho and Amir [16] (see below).

### The peculiar nature of the *D* period

The principal reward of the concurrent cycles framework is is to robustly explain two puzzling trends found in the *C* + *D* period and not accounted by the current literature: (i) *C* + *D* period duration is anticorrelated with single-cell growth rate with a near-inverse pattern and (ii) the amount of growth during this period is anti-correlated with cell size [29].

It is important to spell out how these ingredients lead to problems in existing models. Wallden and coworkers need to fit the duration of the *C* + *D* period to a power law with variable o set and exponent, without any justification. Additionally, they have to assume large mother-daughter correlations in growth rate, and our analysis (Fig. 6 and SI Fig S3) shows that the measured growth-rate correlations do not justify the observed patterns. Additionally while all models can incorporate a size-coupled *C* + *D* period (nonzero *λ*_*C*+*D*_) as an extra ingredient, this ingredient does not have a natural explanation, and we have shown how this choice still leads to problems with the data when the *B* and *C* + *D* period are considered jointly (Fig. 5).

We also stress that the basic ingredient of our model, i.e., concurrence between two synchronous cycles, is different from the hypothesis of parallel subperiods of previous models [16, 34], that basically can be assimilated to the “ICD” framework presented here. In all these models only one replication-related process (the *C* + *D* period) sets division and runs in parallel with another process (the *I* period) setting the inter-initiation time. By contrast, in the concurrent-cycles formalism, cell division can be controlled by the slowest of *two different processes*, only one of which is related to replication.

A notable consequence of concurrent cycles is that the *D* period has a very peculiar status, since its duration is subject to the joint control of both concurrent processes. Consequently, this period is the result of multiple processes including (i) a minimum time from termination to division necessary to complete segregation (at a given growth condition) in case the replication-related process triggers division (ii) the additional “residual” time necessary to wait for the completion of the inter-division process in case this process is the slowest one and determines division. This suggests an “adaptable” duration for the *D* period depending on the size reached at termination [14].

### Size control at replication initiation

Our analysis supports the conclusion that the correlation of initiation size and initial size is not negligible, at least in some data sets. In order to draw this conclusion, we re-analyzed the SeqA data available in the literature, as well as the DnaQ data from Wallden and coworkers. In agreement with these authors, we find that secondary initiations are possible. By performing this analysis we show that, in this data, initiation size may carry some weak but noticeable positive correlation with initial size. In addition, the occurrence of double initiations, on theoretical grounds, is necessary to restore steady conditions from a perturbation. Suppose for example that a cell “misses” a replication initiation event, it will then need a double initiation in order to align its size and chromosome content to the average of its population. Therefore, it seems more plausible that the SeqA behavior is more due to some specific property of this molecule, rather than symptom of a constraint on replication initiation.

Taking this question aside, and assuming that the correlation pattern of the observed points is meaningful, we developed an algorithm to score correlations for clouds of points cut by known constraints, allowing to extract information on the real correlation between volume at initiation and initial volume from the SeqA data sets from ref. [15], removing the spurious correlations arising from the fact that points are available only for the initiation volumes larger than the volume at birth. Also in this case, we detected weak but significant positive correlations.

Both the ICD model and the concurrent cycles model show that the weak correlation observed between initial size and initiation size does not necessarily point to sizer control. Indeed, the same correlation pattern can be found in the scenario of an adder (or a different control) between subsequent initiations. In a framework of concurrent cycles, the reason for this is that the correlation between the two variables ‘initial size’ and ‘initiation size’, is decreased by the possibility that the two different processes decide division.

Overall, comparing the concurrent-cycles model with data, and considering all observed correlation patterns jointly, a scenario of adder control between subsequent initiations appears to be the most plausible (Fig. 7 and SI Fig. S7). It is remarkable that the model can make this prediction, while data analysis cannot settle directly this point. In the future, direct measurements of added volume between subsequent initiations will likely settle this point directly.

### Tuning of the size-scale of the competing circuits

Two additional auxiliary pieces of evidence are important. First, two recent studies measured changes in cell size under several genetic and molecular perturbations affecting key variables such as metabolism, replication, synthesis of essential cell components, cell-cycle proteins etc. [31, 32]. Assuming that initiation triggers cell division after an average constant time, the “*C* + *D*” period, both studies conclude that on average cells initiate replication at a critical size per origin [31, 32], as hypothesized classically by Donachie [3, 5, 6, 35] [36]. Second, cell sizes show “scaling”, and a single size scale defines the probability distribution of initial (at birth) sizes and interdivision times [21, 25]. In other words, the histograms of these variables collapse (across conditions) when rescaled by their means. Together, these two facts indicate that the cell cycle encodes a unique size scale for the cells, and that this scale corresponds to the average size per origin at initiation. Since size is set by the decision to divide, this means that either a single rate-limiting process typically decides cell division [18], or if multiple processes act on similar time scales they need to be tuned in a way that their characteristic size and time scales coincide. Additionally, these processes have to be informed on (or inform) origin number or genome amount.

The concurrent-cycles assumption is meaningful if typically there is competition between the two concurrent processes. This means that in a given condition, cells have non-negligible (and non-small) probability to divide with either of the concurrent processes. In order for this to be the case, the intrinsic size scales of the two processes (mean added size between divisions and mean size at initiation) have to be comparable (and proportional). Any limit where only one process dominates simply reduces to the previously available models [15, 18, 37] and makes our formalism redundant. The matching or near-matching between the two size scales is also a necessary consequence of the existence of a single size scale determining the probability distribution of cell size in *E. coli* [21, 25].

Targeted perturbations of the cell cycle may also support the hypothesis that these scales are matched. For example, deletion of SlmA, the “nucleoid occlusion” protein preventing *E. coli* cells from dividing in presence of unsegregated chromosomes, leaves mean cell size unaffected [10, 38]. We interpret this as a clue in favor of self-tuning of the intrinsic size scale of the inter-division process and the size scale set by replication initiation. Since the two concurrent processes may not compete (or compete less) in these mutants, our prediction is that, looking at the behavior of single cells, the size distribution and the size-timing correlation patterns of these mutants should differ from the wild type.

Finally, competition between concurrent cycles could explain why *E. coli* cells growing at extremely slow growth rates deviate from adder correlations. This could come from variations in the frequency at which each process is bottleneck (*p*_*H*_ in our model). We also observe that even in presence of adder control both between subsequent initiations and between subsequent divisions, the concurrent-cycles model at matched size scales gives a stronger control between division than an adder (in line with data) [39].

## Acknowledgments

We thank Sven van Teeffelen, Clotilde Cadart, Bianca Sclavi, Ilaria Iuliani, Ariel Amir, Po Yi Ho, Nancy Kleckner and Sander Tans, for helpful feedback on this work. This work was supported by the International Human Frontier Science Program Organization, grant HFSP RGY0070/2014. MO was supported by the “Departments of Excellence 2018 - 2022” Grant awarded by the Italian Ministry of Education, University and Research (MIUR) (L. 232/2016). JG was supported by an Omidyar Postdoctoral Fellowship at the Santa Fe Institute. GM was supported by grant nr. 31003A 169978 from the Swiss National Science Foundation to Martin Ackermann.

